# A pillar/perfusion plate enhances cell growth, reproducibility, throughput, and user friendliness in dynamic 3D cell culture

**DOI:** 10.1101/2023.02.16.528892

**Authors:** Vinod Kumar Reddy Lekkala, Soo-Yeon Kang, Jiafeng Liu, Sunil Shrestha, Prabha Acharya, Pranav Joshi, Mona Zolfaghar, Minseong Lee, Paarth Jamdagneya, Sohan Pagnis, Arham Kundi, Samarth Kabbur, Ung Tae Kim, Yong Yang, Moo-Yeal Lee

## Abstract

Static three-dimensional (3D) cell culture has been demonstrated in ultralow attachment well plates, hanging droplet plates, and microtiter well plates with hydrogels or magnetic nanoparticles. Although it is simple, reproducible, and relatively inexpensive, thus potentially used for high-throughput screening, statically cultured 3D cells often suffer from the necrotic core due to limited nutrient and oxygen diffusion and waste removal and have limited *in vivo*-like tissue structure. Here, we overcome these challenges by developing a pillar/perfusion plate platform and demonstrating high-throughput, dynamic 3D cell culture. Cell spheroids have been loaded on the pillar plate with hydrogel by simple sandwiching and encapsulation and cultured dynamically in the perfusion plate on a digital rocker. Unlike traditional microfluidic devices, fast flow rates were maintained within perfusion wells, and the pillar plate could be separated from the perfusion plate for cell-based assays. It was compatible with common lab equipment and allowed cell culture, testing, staining, and imaging *in situ.* The pillar/perfusion plate enhanced cell growth by rapid diffusion, reproducibility, assay throughput, and user friendliness in dynamic 3D cell culture.

## Introduction

Although animal models have been used widely to investigate toxicity and efficacy of drug candidates in preclinical evaluations, the results obtained from animals have not been translated well to humans due to significant difference in genetic makeup^1^. Thus, human cell models have been used widely in drug discovery processes, including high-throughput screening (HTS). The critical problem is that almost all drug candidates identified with two-dimensional (2D) cell monolayers in microtiter well plates in HTS fail when examined in preclinical evaluations^2^. This large failure rate hinges on the fact that 2D cells are physiologically irrelevant to *in vivo.* As an alternative approach, there have been significant advancements made in three-dimensional (3D) cell culture, which include cell spheroids, organoids derived from induced pluripotent stem cells (iPSCs) and adult stem cells (ASDs), 3D-bioprinted tissue constructs, and multi-layered cells in microfluidic devices (a.k.a., organ-on-chips)^3,4^. These 3D cell/tissue models are considered the most appropriate systems for predictive assessment of drug candidates due to more accurate representation of *in vivo* cell characteristics. Nonetheless, there are several technical challenges to adopt 3D cell/tissue models in HTS, which include the necrotic core due to limited nutrient and oxygen diffusion and waste removal, low assay throughput in cell staining and imaging, poor reproducibility in long-term cell culture, and incompatibility with common analytical lab equipment. For example, cell spheroids generated in ultralow attachment (ULA) well plates, hanging droplet plates, and microtiter well plates with hydrogels or magnetic nanoparticles could be used in HTS due to their simplicity, reproducibility, and relatively low costs^3,5,6^. However, cell spheroids cultured in a static condition often suffer from the necrotic core during long-term culture because of the diffusion/transportation issue and have limited *in vivo*-like tissue structure^7^. As compared to cell spheroids, organoids and 3D-bioprinted tissue constructs are more complex and could recapitulate *in vivo* tissue characteristics but still suffer from the necrotic core due to lack of vascular structure^8,9^. To avoid this issue, dynamic cell/tissue culture in petri dishes, spinner flasks, and rotating wall vessels have been used mainly for scale-up production of 3D cells^3,10–12^. The 3D cells cultured in a dynamic condition are often not amenable to HTS systems such as microtiter well plates, including 96-, 384-, and 1536-well plates, due to difficulty in dispensing large cell aggregates robustly and reproducibly using robotic liquid dispensers. Alternatively, microscale dynamic cell culture systems such as microfluidic devices have been used to load, culture, and test cells *in situ.* However, microfluidic devices have inherently low assay throughput due to tedious manual steps necessary for cell loading and media change, incompatibility with HTS equipment due to micropumps necessary to supply growth media, and difficulty in scale-up production and quality control due to complicated steps and highly specialized skills necessary^3,13–15^. To address these challenges, we have developed a pillar/perfusion plate platform, including a 36-pillar plate with sidewalls and slits (36PillarPlate) and a complementary 36-perfusion well plate with reservoirs and microchannels (36PerfusionPlate) *via* injection-molding of polystyrene for dynamic culture of 3D cells and tissues for predictive compound screening. We already developed a 384-pillar plate with a flat pillar surface for microarray 3D bioprinting of cells in hydrogels on the pillars, static culture of cell spheroids, and metabolism-induced hepatotoxicity and neurotoxicity assays in high throughput^16–18^. In the present work, the pillar plate has been further improved to incorporate sidewalls and slits for long-term 3D cell culture, and a complementary perfusion well plate has been developed to reduce cell death in the core and enhance cell growth by dynamic culture. The bidirectional flow rates and velocity profiles within the perfusion wells were simulated with SolidWorks, which were confirmed experimentally by measuring overall flow rates at different liquid volumes and titling angles. Unlike other perfusion well plates^19,20^, the flow rate accelerated under the pillars in the perfusion wells because of the small cross-sectional area. To investigate the effects of the enhanced flow rate under the pillars and improve reproducibility of dynamic 3D cell culture, we have developed a simple spheroid transfer and encapsulation method on the pillar plate with Hep3B cell spheroids generated in a ULA 384-well plate. Overall, the pillar/perfusion plate enhanced cell growth significantly by reducing the necrotic core, reproducibility by robust spheroid transfer, assay throughput by cell staining and imaging *in situ,* and user friendliness without using external pumps and tubes in dynamic 3D cell culture.

## Results

### Design of a 36PillarPlate and a 36PerfusionPlate

Prior to injection molding of the 36PillarPlate and the 36PerfusionPlate, the structure of pillars with sidewalls and slits was designed with SolidWorks, and their function was tested iteratively with 3D printing of prototypes. The final design of the 36PillarPlate with four sidewalls and four slits was determined to improve user friendliness in cell loading and accelerate diffusion in 3D cell culture with fluid flow (**Fig. 1A and 1D**). The 36PillarPlate contains 6 x 6 array of pillars (total 36 pillars per plate) which allows to load up to 6 μL of cell suspension in hydrogels per pillar. Four different methods can be used to load cells in hydrogels on the pillar plate: (1) 3D bioprinting of cell suspension in hydrogels on the pillars^16–18^, (2) manual dispensing of cells with multichannel pipettes, (3) stamping of the pillars into the 384-wells with cell suspension in hydrogels^21^, and (4) transfer of cell spheroids onto the pillars from the ULA 384-wells (presented in **Fig. 3A**). After cell loading on the pillar plate, cells can be encapsulated in hydrogels by thermal gelation *(e.g.,* Matrigel) or ionic crosslinking *(e.g.,* alginate). Cells in hydrogels on the pillar plate can be cultured statically in a 384-deep well plate or dynamically in a perfusion well plate. The complementary 36PerfusionPlate contains 6 x 6 array of perfusion wells (total 36 wells per plate) which are connected by microchannels and reservoirs for dynamic fluid flow (**Fig. 1B and 1E**). The 36PerfusionPlate contains six fluidic channels, each channel consisting of six perfusion wells connected by microchannels and two reservoirs (upper and lower). Therefore, six different conditions can be tested simultaneously with six replicates per condition. The typical volume of cell culture media required in each channel is 600 – 1,000 μL with approximately 60 μL of media in each perfusion well. One of unique features of the pillar/perfusion plate is its flexibility and user friendliness. The pillar plate with 3D-cultured cells in a dynamic condition can be separated from the perfusion well plate and then sandwiched onto the 384-deep well plate for compound testing and cell imaging in a static condition, and *vice versa.* None of existing microfluidic devices provide this kind of flexibility. Due to its small dimension and footprint in a 384-well format, the pillar/perfusion plate is compatible with common analytical equipment such as microtiter well plate readers and bright-field/fluorescence microscopes. Cell growth over time can be monitored easily on the pillar/perfusion plate. By sandwiching the 36PillarPlate with the 36PerfusionPlate (**Fig. 1C and 1F**), cells up to 1.4 mm in diameter can be cultured dynamically on the pillars on a digital rocker in a CO_2_ incubator by generating bidirectional flow.

**Figure 1.**
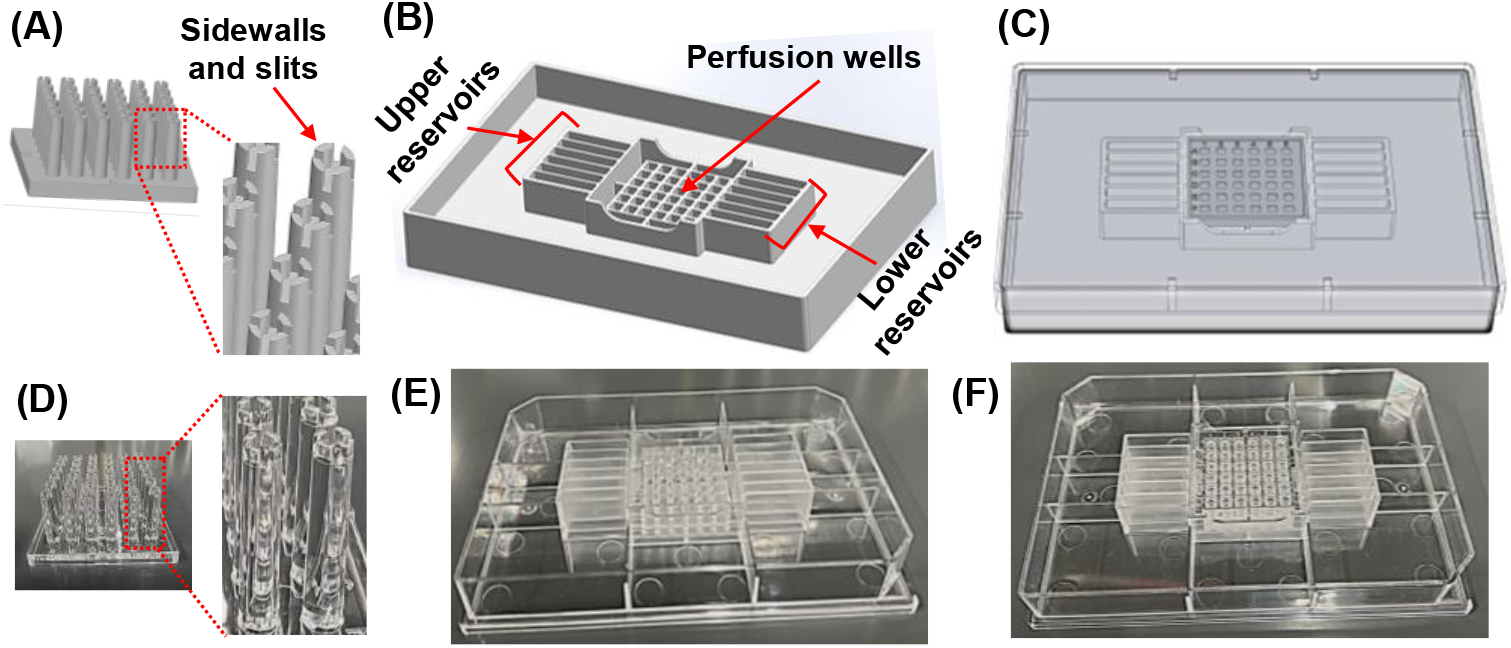
A 36PillarPlate and a complementary 36PerfusionPlate for dynamic 3D cell culture. The SolidWorks design of **(A)** the 36PillarPlate, **(B)** the 36PerfusionPlate, and **(C)** the 36PillarPlate sandwiched onto the 36PerfusionPlate. The picture of **(D)** the 36PillarPlate, **(E)** the 36PerfusionPlate, and **(F)** the sandwiched plates, both manufactured by injection molding with polystyrene.

### Simulation of bidirectional fluid flow within the pillar/perfusion plate

Another unique feature of the pillar/perfusion plate is its rapid flow within the perfusion wells, which could accelerate nutrient and oxygen diffusion and waste removal in dynamic 3D cell culture. To better understand the velocity profile underneath the pillars in the perfusion wells, flow simulation has been performed in the 36PerfusionPlate coupled with the 36PillarPlate by using SolidWorks software. The velocity profile under 10 μm of the pillars in the 36PerfusionPlate was simulated with SolidWorks with 1,000 μL of water per channel at 10° tilting angle and 1 minute frequency of tilting angle change on a digital rocker (**Fig. 2C and Supplementary Video 1 and 2**). Interestingly, fluid flow rates accelerated underneath the pillars in the perfusion wells, which could be explained by Bernoulli’s principle^22^. When a fluid flows within a pipe, the speed of the fluid can be increased if the cross-sectional area of the pipe is reduced. Due to the pillars sandwiched into the perfusion wells, the cross-sectional area of the perfusion wells is reduced significantly, leading to increased flow rates underneath the pillars in the perfusion wells (**Fig. 2B and 2C**). None of perfusion well plates commercially available, including Reprocell’s Alvetex perfusion plate, Lena Bio’s PerfusionPal^19^, and InSphero’s Akura Flow MPS^23^, allow rapid fluid flow within their perfusion wells. In addition, forward and backward flow rates in the 36PerfusionPlate coupled with the 36PillarPlate were simulated with 600, 800, and 1,000 μL of water in each channel at 5°, 10°, and 15° tilting angles and 1 minute frequency of tilting angle change on a digital rocker (**Fig. 2D**). The goal of flow simulation and experimental flow measurement was to find out the optimum operating conditions without overflow of water from the perfusion wells and complete drainage of water in the reservoirs and estimate peak and average flow rates at various water volumes and tilting angles (**Table 1**). From the flow simulation, we quickly noticed that there was no overflow of water from the perfusion wells at 5°, 10°, and 15° tilting angles with 600 μL, 800 μL, and 1,000 μL of water in each channel. In addition, there was complete drainage of water in the upper reservoirs at high tilting angles due to insufficient water in each channel, which was confirmed experimentally. With 1 minute frequency, 600 μL of water in the upper reservoirs was completely drained to the lower reservoirs within 26 ± 1 seconds at 15° tilting angle and 57 ± 3 seconds at 10° tilting angle whereas it took 49 ± 5 seconds at 15° tilting angle with 800 μL of water. Water was not completely drained to the lower reservoirs in the other conditions. Overall, the average flow rates obtained from the experiments were ranged between 2.3 – 7.7 μL/second with increased flow rates with higher tilting angle whereas the average flow rates obtained from the simulation were ranged between 3.3 – 9.3 μL/second (**Table 1**). Thus, the experimental results were comparable to those obtained from the SolidWorks simulation. As compared to typical volumetric flow rates in microfluidic devices (0.008 – 0.3 μL/second), the flow rates in the pillar/perfusion plate are 100 to 1,000-fold faster. Interestingly, the increased tilting angle from 5° to 15° led to significantly increased flow rates whereas the increased water volume in each channel from 600 μL to 1,000 μL minimally influenced the flow rates. This could be because fluid flow in the perfusion plate is generated by gravity from the height difference between the upper and lower reservoirs. The height difference generated by increased tilting angle is much greater than that generated by increased water volume. Based on the results of flow rates from various operating conditions, we decided to use 1,000 μL of cell culture medium in each channel and operate the pillar/perfusion plate on a digital rocker at 10° tilting angle and 1 minute frequency for dynamic 3D cell culture.

**Figure 2.**
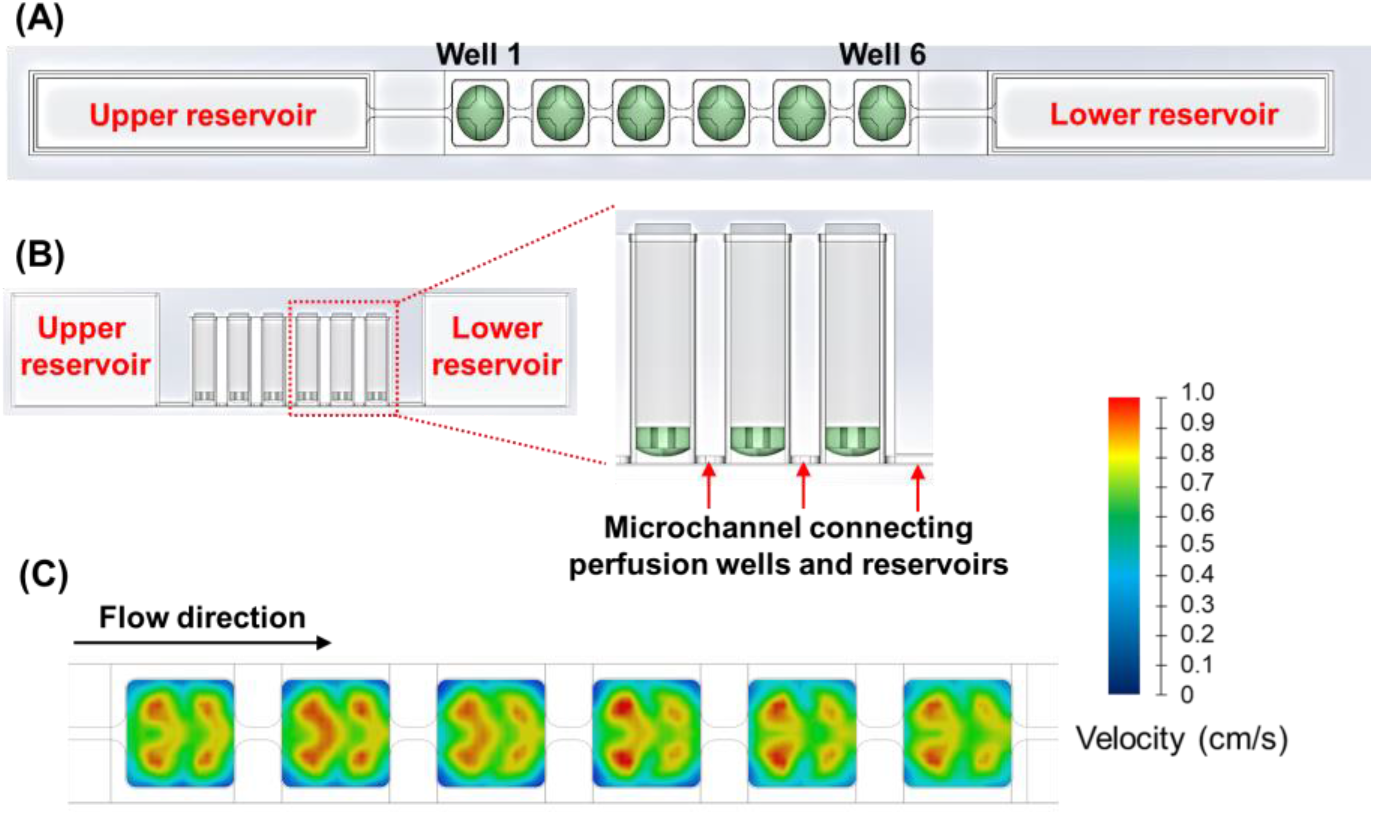
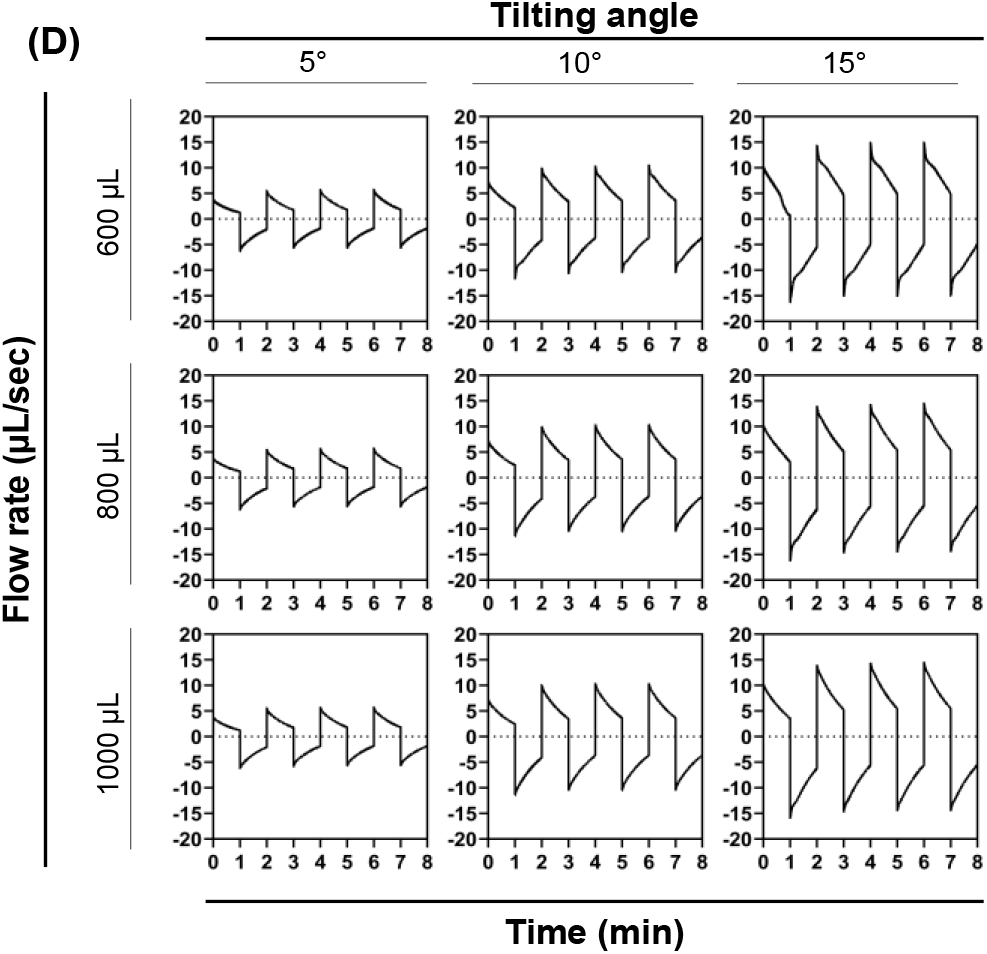
Flow simulation in the 36PerfusionPlate coupled with the 36PillarPlate. **(A)** Top view of the sandwiched plates, illustrating the pillars inserted into the perfusion wells. **(B)** Cross-sectional view of the sandwiched plates. Six perfusion wells and two reservoirs are connected by microchannels. **(C)** Velocity profile under 10 μm of the pillars in the 36PerfusionPlate simulated with SolidWorks with 1,000 μL of water at 10° tilting angle and 1 minute frequency. **(D)** Forward and backward flow rates in the 36PerfusionPlate coupled with the 36PillarPlate simulated with SolidWorks at various water volumes, tilting angles, and 1 minute frequency.

**Table 1.** Average volumetric flow rates obtained from experiment and SolidWorks simulation in the 36PerfusionPlate sandwiched with the 36PillarPlate.

| Total volume | Tilting angle | Average flow rate ($\mu\text{L}/\text{sec}$) obtained from experiment | Average flow rate ($\mu\text{L}/\text{sec}$) obtained from simulation | Peak flow rate ($\mu\text{L}/\text{sec}$) obtained from simulation |
| --- | --- | --- | --- | --- |
| 600 $\mu\text{L}$ | 5° | $2.3 \pm 0.3$ | 3.3 | 5.7 |
| | 10° | $4.0 \pm 0.6$ | 6.3 | 10.4 |
| | 15° | $5.8 \pm 0.8$ | 8.9 | 14.8 |
| 800 $\mu\text{L}$ | 5° | $3.8 \pm 0.4$ | 3.3 | 5.7 |
| | 10° | $4.7 \pm 0.7$ | 6.3 | 10.4 |
| | 15° | $7.0 \pm 0.9$ | 9.2 | 14.4 |
| 1,000 $\mu\text{L}$ | 5° | $4.0 \pm 1.1$ | 3.3 | 5.7 |
| | 10° | $6.4 \pm 0.3$ | 6.4 | 10.4 |
| | 15° | $7.7 \pm 0.4$ | 9.3 | 14.4 |

### Dynamic 3D culture with Hep3B cell spheroids

With the unique features of the pillar/perfusion plate including small cell volume (5 μL) on the pillars and rapid flow underneath the pillars in the perfusion wells, we demonstrated dynamic 3D culture with Hep3B cell spheroids and investigated whether enhanced diffusion due to rapid flow can accelerate cell growth and reduce cell death in the core of spheroids. In addition, we developed a simple spheroid transfer and encapsulation method in alginate on the pillar plate (**Fig. 3A**). Briefly, Hep3B cell spheroids were created in 3 days in a ULA 384-well plate with 5,000 cells seeded in each well. The pillar plate with 5 μL of 1% alginate was sandwiched onto the ULA 384-well plate, and the sandwiched plates were immediately inverted to induce Hep3B cell spheroids settled on the pillars. The cell culture medium in the ULA 384-well plate was supplemented with 2 mM CaCl2 to form partial gelation of alginate during spheroid transfer. After inverting the sandwiched plates back and separating the pillar plate from the ULA 384-well plate, the pillar plate with spheroids was sandwiched onto a 384-deep well plate with 10 mM CaCl2 for complete gelation of alginate for long-term 3D cell culture. The pillar plate was sandwiched onto the perfusion well plate for static and dynamic 3D spheroid culture (**Fig. 3B**). As a control, Hep3B cell spheroids created in the ULA 384-well plate were cultured in a static condition. Overall, the success rate of spheroid transfer and encapsulation on the pillar plate was 95 – 100%. Hep3B cell spheroids were cultured in static and dynamic conditions up to 7 days to monitor changes in spheroid size and cell viability over time. Although the spheroid size reduced slightly after alginate encapsulation on the pillar plate at Day 1 (**Fig. 4A and 4B**), the spheroids grew gradually over time at different rates. Interestingly, the size of spheroids cultured statically in the ULA 384-well plate (I) and dynamically on the pillar plate (III) was almost identical (1,050 μm vs. 1,056 μm after 7 days of culture), but cell viability measured with ATP content using a 3D CellTiter-Glo® assay kit differed nearly two-fold (**Fig. 4C**). It was due to the necrotic core generated in the static culture in the ULA 384-well plate (**Fig. 4E**). From the live/dead cell staining with calcein AM and ethidium homodimer-1 (EthD-1), it was clear that dynamic 3D cell culture on the pillar plate can enhance cell growth and reduce cell death in the core (**Fig. 4E and 4F**). This result was well correlated with the viability measurement with the ATP assay. The coefficient of variation (CV) measured on the pillar plate was below 9% (**Fig. 4D**), indicating the robustness of our approach for cell-based assays. Cell staining and imaging with fluorescent dyes as well as luminescence measurement were performed on the pillar plate *in situ* and measured with a fluorescence microscope and a microtiter well plate reader in the lab, which made our platform highly user friendly and attractive for HTS.

**Figure 3.**
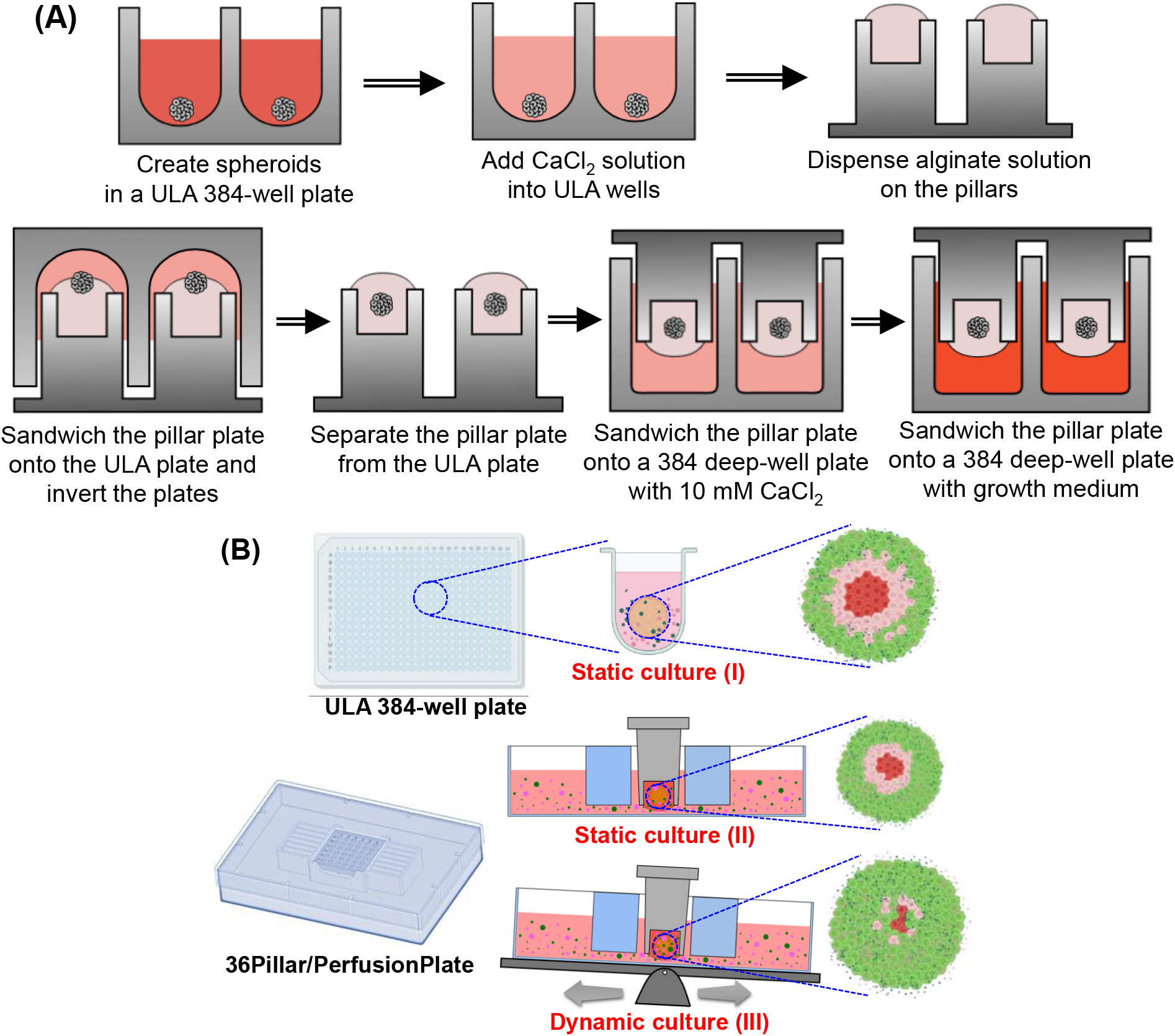
**(A)** Schematic illustration of spheroid transfer from an ultralow attachment (ULA) 384-well plate to the 36PillarPlate. Spheroids formed in the ULA plate were transferred to the 36PillarPlate loaded with alginate. The pillar plate with spheroids was immersed in a 384DeepWellPlate with growth medium containing CaCl2 for spheroid encapsulation and alginate gelation. **(B)** Schematic illustration of Hep3B cell spheroid culture in static and dynamic conditions in the ULA 384-well plate and on the 36PillarPlate sandwiched onto the 36PerfusionPlate. Hep3B cell spheroids were formed in the ULA plate at 5,000 cells/well seeding, which were continuously cultured in the ULA plate in a static condition or transferred to the 36PillarPlate and cultured in static and dynamic conditions in the 36PerfusionPlate. The green dots illustrate live cells whereas the red dots represent dead cells.

**Figure 4.**
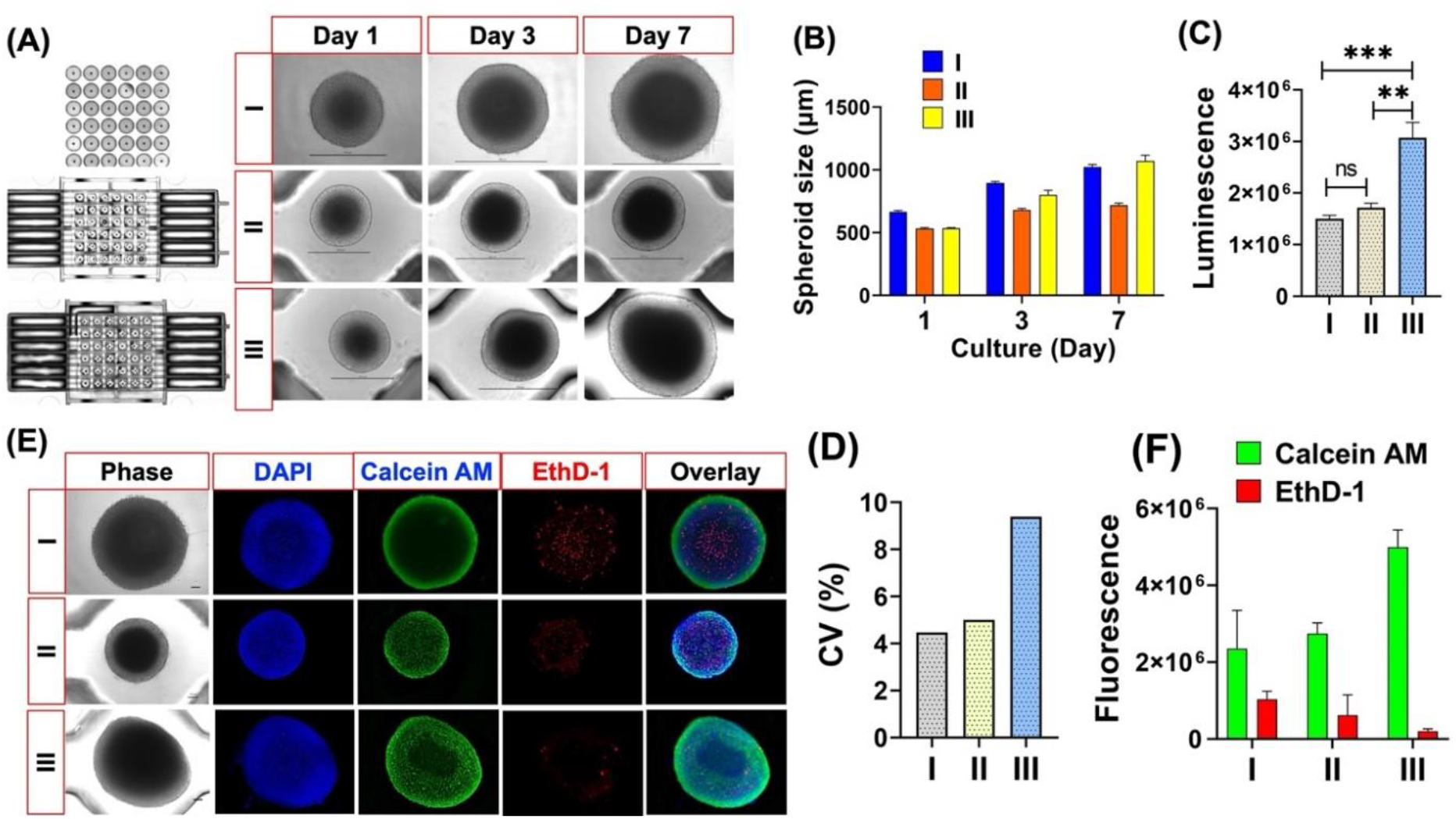
Measurement of spheroid growth rate and cell viability: (I) Static culture in the ULA 384-well plate, (II) static culture on the pillar/perfusion plate, and (III) dynamic culture on the pillar/perfusion plate. **(A, B)** Growth rate of Hep3B cell spheroids over time. The spheroid diameter was measured with 10 spheroids per condition. Scale bar = 500 μm. **(C)** Cell viability measured after 7-day culture with CellTiter-Glo® luminescent cell viability assay kit. n = 12. Student t-test was used to determine the statistical significance. ** = P value 0.001, *** = 0.0001, and ns = non-significant. **(D)** Coefficient of variation (CV) calculated from the cell viability measurement. **(E)** Representative images of live and dead cells in Hep3B cell spheroids after 7-day culture, which were measured with 1 μM calcein AM and 2 μM ethidium homodimer-1 (EthD-1). DAPI was used to stain nuclei. Scale bar = 100 μm. **(F)** Fluorescence intensity obtained from calcein AM and ethidium homodimer-1 stained Hep3B cell spheroids. n = 3.

With the promising results from dynamic 3D cell culture on the pillar/perfusion plate, we investigated the mechanism of cell death in the core of Hep3B spheroids. Potential hypoxia in the core of Hep3B cell spheroids was measured with Image-iT™ green hypoxia reagent (**Fig. 5**). Interestingly, Hep3B cell spheroids were not stained in green with the hypoxia reagent, indicating that cell death in the core could be not because of oxygen depletion but because of insufficient nutrients available. The hypoxia reagent worked normally when Hep3B cell spheroids were incubated in a Ziploc bag to limit oxygen supply. According to the manufacturer instructions^24^, cells should be green fluorescence when the oxygen level in the core is below 5%. Furthermore, Hep3B cell spheroids cultured for 7 days in static and dynamic conditions were assessed with qPCR for the expression levels of anti-apoptotic (cell survival) and pro-apoptotic (cell death) markers (**Fig. 6**). Anti-apoptotic markers such as B-cell lymphoma 2 (BCL2) and myeloid-cell leukemia 1 (MCL1) were significantly upregulated in dynamically cultured Hep3B spheroids on the pillar/perfusion plate whereas pro-apoptotic markers such as BCL2 associated X (BAX) and BCL2 associated agonist of cell death (BAD) were significantly downregulated in both statically and dynamically cultured Hep3B spheroids on the pillar/perfusion plate. Apoptotic cell death is regulated *in vitro* and *in vivo* by the expression levels of anti-apoptotic and pro-apoptotic proteins^25^. It is well known that nutrient depletion causes cell death through BCL2 protein cascade destabilization^26–28^. As the spheroids grow larger, transportation of nutrients into the core could be insufficient, leading to cell death^29^. The enhanced cell growth and reduced cell death on the pillar/perfusion plate indicate the superiority of our platform in dynamic 3D cell culture.

**Figure 5.**
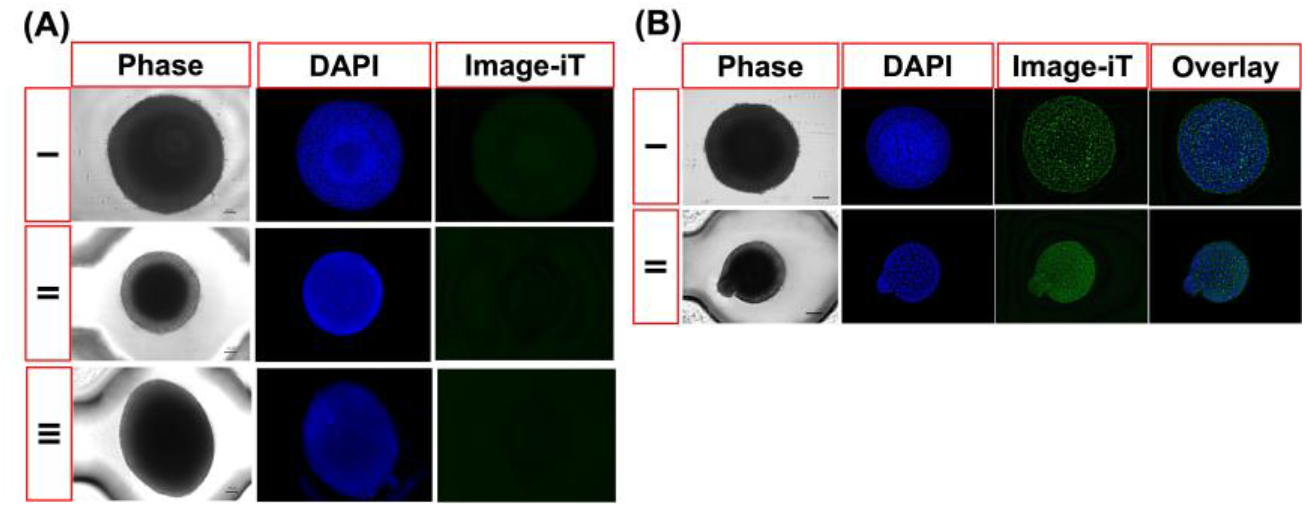
Measurement of hypoxia in the core of Hep3B cell spheroids with Image-iT™ green hypoxia reagent. **(A)** Hep3B cell spheroids cultured for 7 days in a CO_2_ incubator. Scale bar = 100 μm. No hypoxia detected. **(B)** Hep3B cell spheroids cultured for 7 days in a CO_2_ incubator and then incubated in a Ziploc bag for 12 hours to induce hypoxia. Green dots indicate hypoxic cells.

**Figure 6.**
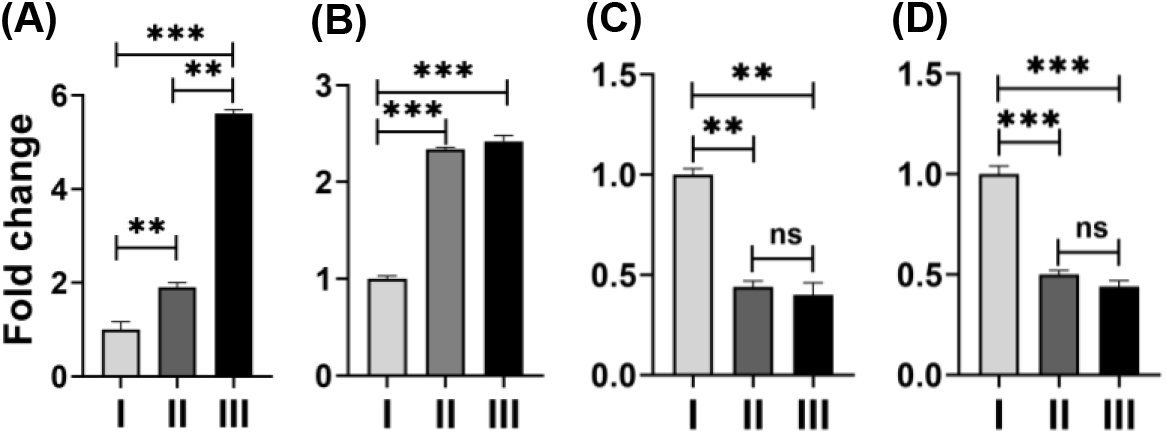
RT-qPCR analysis of anti- and pro-apoptotic gene expression in static and dynamic cultured Hep3B cell spheroids for 7 days: (A) BCL2, (B) MCL1, (C) BAX, and (D) BAD. BCL2 and MCL1 are anti-apoptotic markers whereas BAX and BAD are pro-apoptotic markers. The data was normalized with the expression levels of Hep3B spheroids grown in the ULA plate. A total of 36 spheroids were used to obtain error bars. Student t-test was used for statistical analysis. ** = P value 0.002, *** = 0.0002, and ns = non-significant.

## Discussion

Traditionally, scaffold-free spheroid culture has been performed in ULA plates and hanging droplet plates^3,5,6^. In addition, cell spheroids have been cultured in biomimetic hydrogels to create more complex, miniature tissues such as organoids by mimicking cell-matrix interactions. The critical bottleneck is that spheroid encapsulation in hydrogels is highly labor-intensive because of difficulty in transferring single spheroid with hydrogel to each microtiter well for obtaining reproducible data. Using the pillar/perfusion plate, we demonstrated a simple method of single spheroid transfer and encapsulation in alginate and performed dynamic 3D cell culture with high reproducibility. As compared to traditional 3D cell culture systems, the pillar/perfusion plate has several unique features, which facilitate scale-up production of 3D cells for HTS. First, cell loading on the pillar plate is highly versatile, fast, and reproducible. Unlike extrusion-based 3D bioprinting, microarray 3D bioprinting allows to print hundreds of cell droplets in hydrogels on the pillar plate in a few minutes with the CV values below 25%^16–18^. Second, unlike traditional perfused multiwell plates including Reprocell’s Alvetex perfusion plate, Lena Bio’s PerfusionPal, and InSphero’s Akura Flow MPS, flow rates within the perfusion wells in the 36PerfusionPlate are fast, which enhanced cell growth and reduced cell death in the core of 3D-cultured cells by rapid diffusion of nutrient and oxygen and removal of wastes (**Fig. 4**). In addition, the pillar/perfusion plate allowed to control flow velocity by simply changing the tilting angle of the perfusion plate on a digital rocker without using pumps and tubes. The average linear flow rates in the pillar/perfusion plate (1.5 – 2.0 cm/sec) were comparable with the blood flow rates in veins. For example, the average arterial blood flow rates in the heart are in the range of 10 – 60 cm/sec^30,31^ whereas the average venous blood flow rates in fingers and legs are in the range of 1.5 – 10 cm/sec^32–34^. The average blood flow rates in the capillaries are in the range of 0.03 – 0.3 cm/sec^35–37^. On the other hand, typical linear flow rates in microfluidic devices are in the range of 0.002 – 0.016 cm/sec^38–42^, which are 100 – 1,000-fold slower than those obtained from the pillar/perfusion plate. The slow flow rates in microfluidic devices may be designed to match the blood flow rates in the capillaries or be necessary due to high sheer stress in narrow microchannels where cells are growing. Third, the assay volume on the pillar/perfusion plate is small, thus amenable to high-throughput compound screening. The pillar/perfusion plate is built on the footprint of standard 384-well plates, requiring 5 μL of cells in hydrogels on each pillar (200 – 5,000 cells/pillar) and 50 – 80 μL of culture media in each perfusion well. As compared to conventional 3D culture systems including 6-/24-well plates, petri dishes, and spinner flasks, the assay volume is reduced approximately 20 – 100-fold (**Supplementary Table 1 and 2**). Forth, the pillar/perfusion plate is highly flexible and user friendly for cell culture and imaging. Unlike traditional microfluidic devices, the pillar plate with 3D-cultured cells can be detached from the perfusion well plate or the deep well plate and then sandwiched onto fresh well plates for culture medium changing and high-content imaging (HCI) of cells. Cell image acquisition from 3D-cultured cells is easy and straightforward because the whole sample depth fits within the focus depth of a normal objective (4x and 10x). The amount of cell samples and culture media obtained from the pillar/perfusion plate is sufficient for qPCR and ELISA analysis, which is often a limitation of microfluidic devices. Fifth, the pillar/perfusion plate is compatible with standard 384-well plates and HTS equipment such as automated fluorescence microscopes and microtiter well plate readers, and thus can be easily adopted in current drug discovery efforts. There is no expensive, proprietary equipment necessary to operate the pillar/perfusion plate. The vast majority of current microfluidic devices are inherently low throughput in dynamic 3D cell culture. Except for the OrganoPlate® from MIMETAS, only one cell/tissue culture per device can be performed. Thus, it is urgently necessary to develop a new microfluidic device that can accommodate dynamic 3D cell culture and compound test in parallel massively. The 36PillarPlate coupled with the 36PerfusionPlate allows to culture 3D cells dynamically on 36 pillars, which can be separated and tested individually with 36 compounds in a deep well plate. Other microfluidic devices could not provide such flexibility and user friendliness.

## Outlook

The pillar/perfusion plate platform offers several distinctive advantages over conventional 3D cell culture systems and microfluidic devices, including (1) extremely fast and robust cell loading by microarray 3D printing or spheroid transfer, (2) high flow rates in the center of perfusion wells, reducing cell death in the core of 3D cells, (3) small assay volume for HTS, (4) high compatibility with standard 384-well plates and existing lab equipment, and (5) highly flexible and user-friendly operation without expensive, proprietary equipment. We have successfully demonstrated dynamic culture of Hep3B cell spheroids on the pillar/perfusion plate for scale-up production and potential compound testing. In addition to the 36PillarPlate and the 36PerfusionPlate, we have developed a high-density 144PillarPlate and a complementary 144PerfusionPlate *via* injection-molding of polystyrene for high-throughput dynamic 3D cell culture and HTS. It is worth noting that encapsulation of pluripotent stem cells in biomimetic hydrogels, such as Matrigel, is essential for organoid differentiation and maturation. The pillar/perfusion plate could be used for dynamic human organoid culture to improve throughput and reproducibility in organoid differentiation and maturation. Current 3D cell culture systems are not amenable to organoid printing and culture in hydrogels for high-throughput compound screening. In addition, the pillar/perfusion plate could be used for multi-organ interaction studies as different tissue modules can be cultured on the pillars in each perfusion channel and be combined easily by separating the pillar plate, rotating 90° angle, and sandwiching back into the perfusion plate. The “killer application” of the pillar/perfusion plate is robust, scale-up production of dynamically cultured 3D cells in hydrogels for predictive compound screening.

## Methods

### Preparation of the 36PillarPlate and the 36PerfusionPlate

The 36PillarPlate and the 36PerfusionPlate were manufactured by injection molding with polystyrene and functionalized with hydrophilic polymers (Bioprinting Laboratories Inc., TX, USA). The 36PillarPlate contains 6 x 6 array of pillars in 4.5 mm pillar-to-pillar distance whereas the 36PerfusionPlate contains six fluidic channels, each channel consisting of six perfusion wells in 4.5 mm well-to-well distance connected by microchannels between two reservoirs (**Fig. 1**).

### Flow simulation with SolidWorks

The velocity profile under 10 μm of the pillars in the 36PerfusionPlate was simulated using SolidWorks software package (SolidWorks Research Standard and Simulation Premium 2022, MA, USA) with 600, 800, and 1,000 μL of water at 5°, 10°, and 15° tilting angle and 1 minute frequency of tilting angle change on a digital rocker. In addition, bidirectional flow rates inside the microchannels within the 36PerfusionPlate coupled with the 36PillarPlate were also simulated with SolidWorks. The geometry of the 36PerfusionPlate was simplified to model only one fluid channel consisting of six perfusion wells with two reservoirs in a single row connected by microchannels (**Fig. 2A**). The microchannels have a dimension of 0.4 mm x 0.4 mm. The model assumed that the 36PillarPlate is sandwiched onto the 36PerfusionPlate. There is a 0.5 mm gap from the bottom of perfusion wells to the top of pillars. It was also assumed that the water-air boundary at the top of the perfusion wells acted as a wall. The surface of the perfusion plate was set as polystyrene with a roughness of 1.7 nm. The physical property was directly generated from SolidWorks database by setting the water temperature at 310.15K. The volume flow rate, mass flow rate, and velocity of fluid flow were simulated and recorded using SolidWorks Flow Simulation by numerically solving the Favre-averaged Navier-Stokes equations^43^. To predict the flow dynamics within the 36PerfusionPlate coupled with the 36PillarPlate, SolidWorks Flow Simulation employed transport equations for the turbulent kinetic energy and its dissipation rate (**Fig. 2C and Supplementary Video 1**)^43^.

### Measurement of average flow rates in the 36PerfusionPlate coupled with the 36PilllarPlate

The rates of fluid flow were measured in the 36PerfusionPlate sandwiched with the 36PilllarPlate using phosphate-buffered saline (PBS, Gibco, 10010-023) on a digital rocker (OrganoFlow® L, Mimetas) at 5°, 10°, and 15° tilting angles and 1 minute frequency of tilting angle change. Three different volumes (600 μL, 800 μL, and 1,000 μL) of PBS were added to each upper reservoir of the 36PerfusionPlate, which was kept on a flat surface for 10 minutes to ensure fluid flow evenly through microchannels and reach equilibrium. After sandwiching the 36PilllarPlate onto the 36PerfusionPlate, the sandwiched plates were mounted on the digital rocker at 0° tilting angle. All PBS in the lower reservoir was pipetted out and added to the upper reservoir, and then the tilting angle of the digital rocker was changed abruptly to 5°, 10°, or 15°. After 30 seconds or 1 minute later, the amount of PBS in the upper and lower reservoirs was measured to calculate the volume of PBS flow through the perfusion wells.

### Formation of Hep3B cell spheroids in a ULA 384-well plate

Hep3B human hepatoma cell line (ATCC) was cultured in RPMI 1640 (Gibco, 11875-093) supplemented with 10% fetal bovine serum (Gibco, 26140079) and 1% penicillin-streptomycin (Gibco,15140122). At around 70% confluency, the cells were detached with Trypsin and seeded in an ultra-low attachment (ULA) 384-well plate (Nexcelom Bioscience LLC., USA, ULA-384U) with 5,000 cells seeded in each well. The cell culture was maintained with alternative day medium changes in a humidified 5% CO_2_ incubator at 37°C for 3 days.

### Transfer of Hep3B cell spheroids from the ULA 384-well plate to the 36PillarPlate

Prior to spheroid transfer, 5 μL of 1% (w/v) alginate prepared by dissolving alginic acid sodium salt (Sigma-Aldrich, A1112) in distilled water was dispensed on top of each pillar using a multichannel pipette. The pillar plate with 5 μL of 1% alginate was sandwiched onto the ULA 384-well plate with Hep3B cell spheroids in RPMI 1640 containing 2 mM CaCl2 for partial gelation of alginate. The sandwiched plates were immediately inverted upside down and incubated for 30 minutes in a 5% CO_2_ incubator at 37°C to ensure Hep3B cell spheroids settled on the pillars. After inverting the sandwiched plates back and separating the pillar plate from the ULA 384-well plate, the pillar plate with spheroids in alginate was sandwiched onto a clear-bottom, 384-deep well plate (Bioprinting Laboratories Inc., Texas, USA) with 80 μL of RPMI 1640 containing 10 mM CaCl2 for 10 minutes for complete gelation of alginate for long-term 3D cell culture (**Fig. 3**). The pillar plate was separated and sandwiched onto the perfusion well plate for static and dynamic 3D spheroid cultures for 7 days on the digital rocker in a humidified 5% CO_2_ incubator at 37°C to monitor changes in spheroid size and cell viability over time. As a control, Hep3B cell spheroids created in the ULA 384-well plate were cultured in a static condition for 7 days.

### Measurement of Hep3B spheroid viability

Due to the large size of Hep3B cell spheroids, viability was assessed by measuring the ATP content and the cell membrane integrity. The total ATP contents in static and dynamic cultured Hep3B cell spheroids were measured with the CellTiter-Glo® assay kit (Promega, G9681). Briefly, an equal volume mixture of the CellTiter-Glo reagent and RPMI 1640 was added to the 384-deep well plate (80 μL/well). The 36PillarPlate with Hep3B cell spheroids was sandwiched onto the 384-deep well plate with the CellTiter-Glo reagent and RPMI 1640 for 45 minutes at room temperature. For the ULA 384-well plate with Hep3B cell spheroids, the CellTiter-Glo reagent was added directly into the wells and incubated for 45 minutes at room temperature. The luminescence signal which represents the amount of ATP in viable cells was measured with a microtiter well plate reader (BioTek Cytation 5, Agilent).

The live and dead cells in static and dynamic cultured Hep3B cell spheroids were measured with Live/Dead™ viability/cytotoxicity kit (ThermoFisher, L3224) containing calcein AM and ethidium homodimer-1 (EthD-1). Briefly, RPMI 1640 containing 2 μM calcein AM and 4 μM EthD-1 along with 1 mg/mL DAPI (ThermoFisher, 62248) was added to the 384-deep well plate (80 μL/well). The 36PillarPlate with Hep3B cell spheroids was sandwiched onto the 384-deep well plate with the live/dead cell staining dyes for 2 hours in a humidified 5% CO_2_ incubator at 37°C. For the ULA 384-well plate with Hep3B cell spheroids, the live/dead cell staining dyes were added directly into the wells and incubated for 2 hours in a humidified 5% CO_2_ incubator at 37°C. After staining live cells in green and dead cells in red, the spheroids on the pillar plate and in the ULA 384-well plate were rinsed carefully with PBS and visualized under an automated fluorescence microscope (BZ-X810, Keyence). The mean fluorescence intensity was measured by subtracting the background intensity with ImageJ.

### Measurement of hypoxia

Hypoxia in static and dynamic cultured Hep3B cell spheroids was measured with Image-iT™ reagent (ThermoFisher, I14833). Briefly, RPMI 1640 containing 8 μM Image-iT reagent was added to the 384-deep well plate (80 μL/well). The 36PillarPlate with Hep3B cell spheroids was sandwiched onto the 384-deep well plate with the Image-iT reagent for overnight in a humidified 5% CO_2_ incubator at 37°C. For the ULA 384-well plate with Hep3B cell spheroids, Image-iT reagent was added directly into the wells and incubated for overnight in a humidified 5% CO_2_ incubator at 37°C. On the next day, 1 mg/mL DAPI in RPMI 1640 was added and incubated for 30 minutes. After cell staining, the spheroids on the pillar plate and in the ULA 384-well plate were rinsed carefully with PBS and visualized under the Keyence fluorescence microscope. To induce hypoxia intentionally, the pillar plate and the ULA 384-well plate with Hep3B cell spheroids were sealed in a Ziploc bag and kept in a humidified 5% CO_2_ incubator at 37°C for 12 hours.

### Measurement of anti- and pro-apoptotic marker expression

Since depletion of nutrients causes apoptotic cell death, the expression levels of anti-apoptotic and pro-apoptotic markers in static and dynamic cultured Hep3B cell spheroids were measured by qPCR. Briefly, total RNA was isolated from Hep3B cell spheroids on the pillar plate and in the ULA plate using a RNeasy Plus kit (Qiagen,74134). One μg of RNA was converted into cDNA using the high-capacity cDNA reverse transcription kit according to the manufacturer instructions (Applied Biosystems, 4368814). qPCR was performed with PowerTrack SYBR green master mix (A46109) by using QuantStudio™ 5 qPCR system (ThermoFisher, A34322) with specific primers (**Table 2**). Analysis was done with the ΔΔCt method. The relative gene expression levels were normalized with the housekeeping gene GAPDH.

**Table 2.**
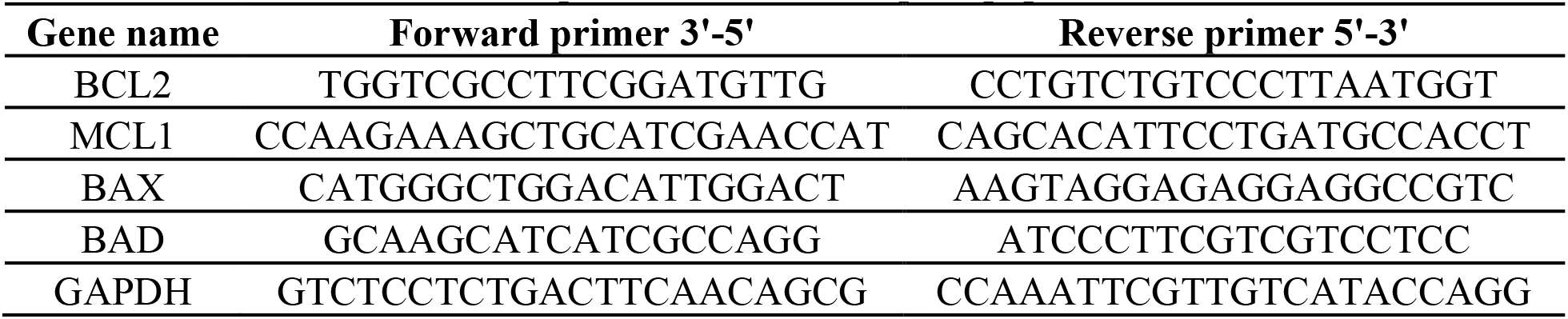
List of primers for anti- and pro-apoptotic markers.

### Statistical analysis

Statistical analysis was done by using GraphPad Prism. Student t-test was performed to determine statistical significance and P values. Data represent mean ± standard deviation.

## Data availability

The authors declare that all the data supporting the results in this study are available within the article and its Supplementary Information.

## Acknowledgments

This study was supported by the National Institutes of Health (NIDDK UH3DK119982, NCATS R44TR003491, and NIEHS R01ES025779) and the Ohio Third Frontier Commission (TVSF Phase IB and Phase II).

## Author contributions

V.K.R.L.: Designed and conducted fluid flow and cell culture experiments, interpreted and analyzed the results, and wrote the manuscript.

S.Y.K.: Tested the injection-molded pillar/perfusion plate, developed surface coating methods, and performed flow simulation with SolidWorks.

J.L, U.T.K., and Y.Y.: Performed flow simulation with SolidWorks and wrote the SolidWorks simulation method section.

S.S.: Developed surface coating methods, performed the spheroid transfer, wrote the spheroid transfer method section, and made schematics for spheroid transfer.

P.A.: Developed surface coating methods, performed the spheroid transfer, and wrote the spheroid transfer method section.

P.J.: Developed surface coating methods and data acquisition.

M.Z.: Helped in handling experiments and writing the introduction section.

M.S.L.: Helped to write the discussion section.

P.J., S.P., A.K., and S.K.: Helped in manuscript editing and created the animation video of dynamic culture.

M.Y.L.: Conceived and designed the pillar/perfusion well plate, developed surface chemistry, planned flow simulation and cell culture experiments, interpreted the results, wrote and revised the manuscript, and supervised the project.

