## Supplementary information for "A pillar/perfusion plate enhances cell growth, reproducibility, throughput, and user friendliness in dynamic 3D cell culture"

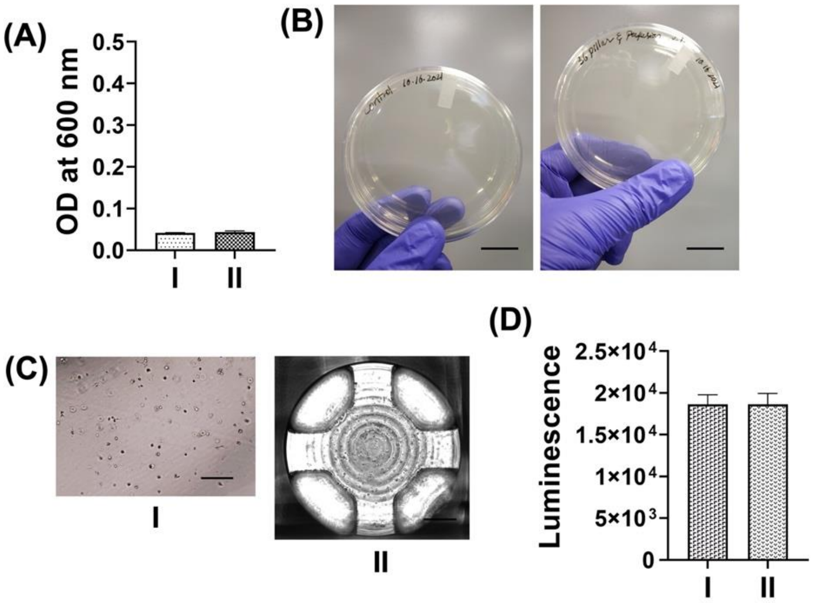


**Supplementary Figure 1. Sterility and biocompatibility tests with (I) a 96-well plate and (II) the 36PillarPlate sandwiched with the 36PerfusionPlate. (A)** Measurement of optical density (OD) at 600 nm with growth media exposed to (I) and (II) for 24 hours to determine microbial contamination. n = 6. **(B)** Measurement of microbial contamination with growth media poured on agar plates. No microbial contamination detected. **(C)** Hep3B cells loaded on (I) and (II) to measure biocompatibility. **(D)** Biocompatibility measured with CellTiter-Glo® 3D assay kit. n = 6. There was no cell death detected.

**Supplementary Table 1.** Comparison of culture medium usage between the 36PillarPlate and traditional microtiter well plates for static cell culture.

| Microtiter well plates | Working volume required (μL) | Working volume for the 36PillarPlate (μL) | Difference in medium usage (fold) |
|---|---|---|---|
| 6-well | 3,000 | | 38 |
| 12-well | 1,500 | | 19 |
| 24-well | 1,000 | 80 | 13 |
| 48-well | 500 | | 6 |
| 96-well | 200 | | 3 |

**Supplementary Table 2.** Comparison of culture medium usage between the 36PerfusionPlate and commercially available perfusion systems for dynamic cell culture.

| Lena Bio perfusion plate | Working volume required (μL) | Working volume for the 36PerfusionPlate (μL) | Difference in medium usage (fold) |
|---|---|---|---|
| 12-well | 3,000 | 80 | 38 |
| 48-well | 1,000 | | 12 |

Culture medium volume in perfusion wells is compared.

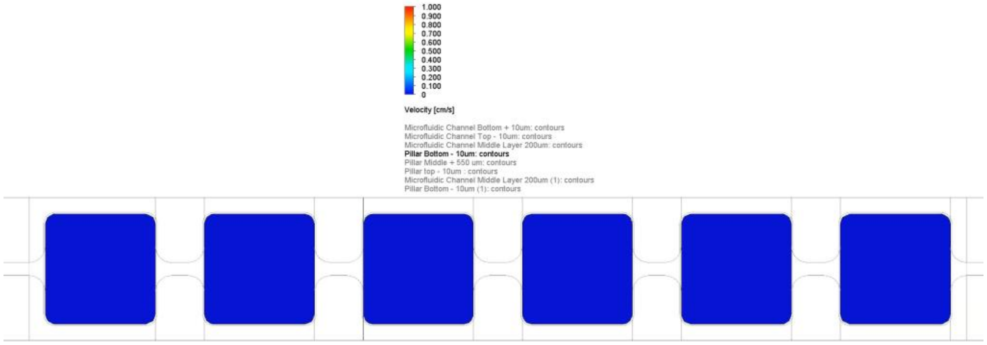


**Supplementary Video 1.** The velocity profile under 10 μm of the pillars in the 36PerfusionPlate simulated with SolidWorks with 1,000 μL of water at 10° tilting angle and 1 minute frequency of tilting angle change (https://youtu.be/nokDLWiRmew). The surface roughness of the 36PerfusionPlate was equal to a petri dish (0.0017 μm).

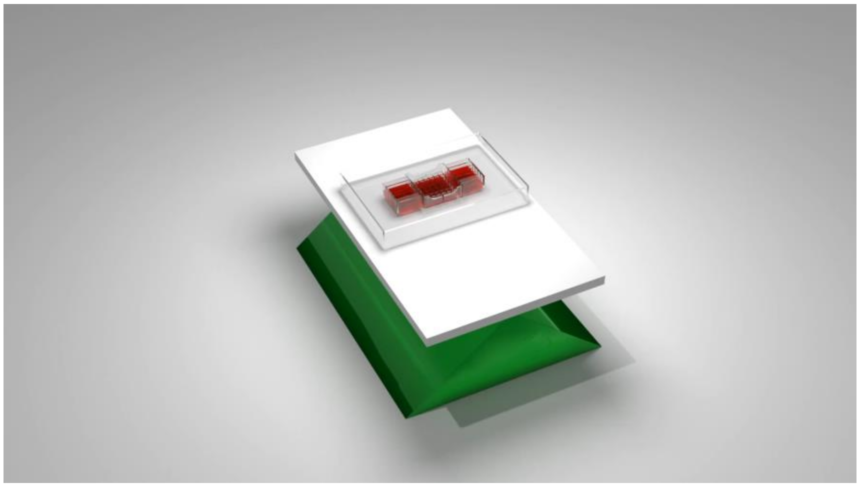

**Supplementary Video 2.** Animation of dynamic spheroid culture on the pillar/perfusion plate on a digital rocker (https://www.youtube.com/watch?v=EQipYdUdOEI).
